# Columnar scale lesions in barrel cortex degrade tactile discrimination but not detection

**DOI:** 10.1101/2022.06.16.496452

**Authors:** Lauren Ryan, Maya Laughton, Andrew Sun-Yan, Ravi Pancholi, Simon Peron

## Abstract

Primary sensory cortices typically display functional topography, suggesting that even small cortical volumes may underpin perception of specific stimuli. Because traditional loss-of-function approaches have a relatively large radius of effect (>1 mm), the behavioral necessity of smaller cortical volumes remains unclear. In the mouse primary vibrissal somatosensory cortex (vS1), ‘barrels’ with a radius of ∼150 μm receive input predominantly from a single whisker, partitioning vS1 into a topographic map of well-defined columns. Here, we train animals implanted with a cranial window over vS1 to perform single-whisker perceptual tasks. We then use high-power laser exposure centered on the barrel representing the spared whisker to produce lesions with an average volume of ∼2 barrels. These columnar scale lesions impair performance on object location discrimination tasks without disrupting vibrissal kinematics. Animals with degraded discrimination performance can immediately perform a detection task with high accuracy. Animals trained *de novo* on both simple and complex detection tasks showed no behavioral deficits following columnar scale lesions. Thus, vS1 barrels are necessary for performing object location discrimination but not simple or complex object detection behaviors.

## Introduction

To understand the neural basis of perception, it is important to identify brain areas that causally contribute to behavior. The behavioral necessity of a brain area is typically assessed by performing loss-of-function perturbations. Across sensory modalities, such experiments suggest that primary sensory cortices are essential for some tasks but not others (Lashley, 1931a, b; Schneider, 1969; Hutson and Masterton, 1986; Newsome and Pare, 1988; Prusky and Douglas, 2004; Glickfeld et al., 2013; Shih et al., 2013; Poort et al., 2015; Goard et al., 2016; Resulaj et al., 2018; Stuttgen and Schwarz, 2018; Slonina et al., 2022). Traditional loss-of-function perturbations, however, have poor spatial resolution: both transient inactivation (O’Connor et al., 2010; Poort et al., 2015; Goard et al., 2016; Li et al., 2019) and permanent lesions (Lashley, 1931a; Hutson and Masterton, 1986; Resulaj et al., 2018) usually have a radius of effect exceeding 1 mm, thereby impacting hundreds of thousands of neurons. In addition, transient perturbations often produce off-target effects that can degrade behavior even in structures whose permanent removal does not perturb behavior. (Otchy et al., 2015; Hong et al., 2018; Wolff and Olveczky, 2018). Thus, more spatially focused and permanent loss-of-function approaches are essential for identifying the minimal subset of cortical activity needed for perception.

In the mouse vibrissal system, individual whiskers project to small, defined patches of primary vibrissal somatosensory cortex (vS1) called ‘barrels’ (radius: ∼150 μm; (Lefort et al., 2009)). Most inactivation studies in vS1 do not explore the behavioral contribution of individual barrels due to poor spatial resolution. For instance, optogenetic inactivation experiments targeting vS1 often directly perturb activity not just in vS1 but also in adjacent structures such as secondary vibrissal somatosensory cortex (Guo et al., 2014a; Li et al., 2019). Lesions of vS1 are also usually extensive, often impacting additional cortical areas along with subcortical structures (Hutson and Masterton, 1986; Hong et al., 2018). Such area-scale perturbations of vS1 degrade performance on aperture size discrimination (Krupa et al., 2001), object location discrimination (O’Connor et al., 2010; Guo et al., 2014b), gap crossing (Shih et al., 2013), and texture discrimination tasks (Guic-Robles et al., 1992). In contrast, object detection is sensitive only to transient inactivation of vS1, as mice can quickly relearn a detection task following vS1 aspiration (Hong et al., 2018). While vS1 is thus implicated in many tasks, it remains unclear whether perceptually relevant activity is mostly confined to the barrels of specific whiskers or is more broadly distributed across vS1.

Here, we precisely lesion small volumes of cortex in awake mice previously implanted with a cranial window using a femtosecond laser. Our approach obviates the need for post-operative recovery and allows for the lesioning of a targeted barrel along with partial removal of adjacent barrels. In mice performing a vibrissal go/no-go object location discrimination task with a single whisker, these lesions, centered around the barrel representing that whisker, degraded performance. Lesions did not impact whisking kinematics. Mice could accurately perform a go/no-go object detection task immediately after a columnar scale lesion in vS1, suggesting that such tasks are not dependent on individual vS1 barrels. Mice trained on a more complex detection task with two response contingencies and a delay period between stimulus presentation and response were also unimpacted by columnar scale lesions. Thus, barrel columns are necessary for object location discrimination but not detection tasks.

## Materials and Methods

### Animals

Adult C57/BL6J (JAX X 000664) mice (5 female, 9 male) and Ai162 (JAX 031562) X Slc17a7-Cre (JAX X 023527) (Daigle et al., 2018) mice (4 female, 3 male) were used (**Table 1**). In cortex, Ai162 X Slc17a7-Cre mice express GCaMP6s in excitatory neurons. To suppress transgene expression during development, Ai162 X Slc17a7-Cre breeders were fed a diet that included doxycycline (625 mg/kg doxycycline; Teklad), so that mice received doxycycline until weaning. All animal procedures were approved by New York University’s University Animal Welfare Committee.

**Table 1.** List of animals. Mice are divided into task-based cohorts. Symbols for individual mice are unique and consistent throughout the manuscript within a cohort. Notes: *, mice with large lesions, excluded for lesion analysis but included in pre-lesion analyses; **, mice with no whisker videography, excluded from lesion and video-dependent analyses; ***, mouse that showed unusually high ignore rate post-lesion, so excluded from post-trim analysis ; †, lesion size estimated due to missing slices (Methods).

| Animal ID | Genotype | Task | Sex | Lesion volume (mm<sup>3</sup>) | Whisker | Behavior start age (weeks) |
| --- | --- | --- | --- | --- | --- | --- |
| 014712* | C57/BL6 | go/no go discrim | F | 1.3214 | C2 | 11 |
| 014377 | C57/BL6 | go/no go discrim | M | 0.1891 | C2 | 9 |
| 014390 | C57/BL6 | go/no go discrim | M | 0.2256 | C2 | 8 |
| 014389 | C57/BL6 | go/no go discrim | M | 0.1483 | C2 | 9 |
| 014388 | C57/BL6 | go/no go discrim | M | 0.0494 | C2 | 9 |
| 014378* | C57/BL6 | go/no go discrim | F | 1.0068 | C3 | 9 |
| 009848 | C57/BL6 | go/no go discrim | F | 0.3156 | C2 | 16 |
| 009840 | C57/BL6 | go/no go discrim | M | 0.162 | C2 | 10 |
| 012266 | C57/BL6 | go/no go discrim | M | 0.0996 | C2 | 9 |
| 296116** | slc17a7-Cre X Ai162 | go/no go discrim | M | 0.1717 | C2 | 15 |
| 283529 | slc17a7-Cre X Ai162 | go/no go detection | M | 0.9621 | C2 | 12 |
| 283541 | slc17a7-Cre X Ai162 | go/no go detection | F | 0.5002 | C2 | 12 |
| 293169 | slc17a7-Cre X Ai162 | go/no go detection | F | 0.5008 | C3 | 16 |
| 293317 | slc17a7-Cre X Ai162 | go/no go detection | F | 0.1405 | C2 | 16 |
| 293158 | slc17a7-Cre X Ai162 | go/no go detection | M | 0.2883 | C2 | 18 |
| 295141*** | slc17a7-Cre X Ai162 | go/no go detection | F | 0.7130† | C3 | 9 |
| 014386 | C57/BL6 | 2 lickport detection | M | 0.0655† | C2 | 21 |
| 014174 | C57/BL6 | 2 lickport detection | F | 1.3214 | C2 | 11 |
| 014173 | C57/BL6 | 2 lickport detection | M | 0.0321 | C2 | 11 |
| 014385 | C57/BL6 | 2 lickport detection | M | 0.1757† | C3 | 11 |
| 014374 | C57/BL6 | 2 lickport detection | F | 0.1286 | C2 | 11 |

### Surgery

Mice (6-10 weeks old) were anesthetized with isoflurane during cranial window and headbar implantation (3% induction, 1.5% maintenance). A titanium headbar was attached to the skull with cyanoacrylate (Vetbond). A circular craniotomy (3.5 mm diameter) was made in the left hemisphere over vS1 (center: 3.5 mm lateral, 1.5 mm posterior from bregma) using a dental drill (Midwest Tradition, FG 1/4 drill bit). Following the craniotomy, a double-layer cranial window (4.5 mm external diameter, 3.5 mm inner diameter; #1.5 coverslip; adhered with Norland 61 UV glue) was placed over the craniotomy. The cranial window and headbar were affixed to the skull with dental acrylic (Orthojet, Lang Dental).

### Behavior

Following surgical recovery, mice were water restricted and trimmed to whiskers C1-3. Mice were trained on one of three tasks. (1) Go/no-go object location discrimination task. A metal pole (0.5 mm diameter; Drummund Scientific, PA, USA) entered into the mouse’s whisking plane in a series of proximal positions or in a series of distal positions. Mice were rewarded with water for licking while the pole was in reach if the pole appeared in one of the proximal positions (in reach time, 2-3 seconds). Licking when the pole appeared in one of the distal positions led to a timeout period and an aversive sound. (2) Go/no-go object detection task. A metal pole entered into the mouse’s whisking plane in or out of reach of its one spared whisker. Mice were rewarded for licking when the pole was in reach and given a timeout period and an aversive sound for licking when the pole was out of reach. (3) Two lickport object detection task. Each trial consisted of three epochs: stimulus epoch (2 s), delay (300 ms), and response (<2 s). During the stimulus epoch, a metal pole entered the mouse’s whisking plane at an in reach proximal position or an out of reach distal position and then moved outside the whisking plane. During the delay epoch, the lickport moved into reach. During the response epoch, an auditory cue indicated the mouse should respond, and mice were rewarded for licking the right lickport during in reach trials, and the left lickport during out of reach trials. Incorrect responses resulted in a timeout period and immediate withdrawal of the lickport. Mice were always trained to criterion (d-prime exceeded 1.5 for two consecutive days) on the task using C1-3 and then trimmed to a single whisker.

All behavioral training proceeded in a standard sequence. First, mice were handled and habituated to the behavioral apparatus. After habituation, mice were trained to lick for a water reward and then proceeded directly to behavioral training on the go/no-go tasks. For the two lickport task, mice also learned the timing of the task and to lick both left and right lickports before training progressed. In the location discrimination task, mice started with distal positions out of reach. Distal positions were quickly brought into reach once mice became accustomed to the task structure and whisked vigorously. In detection tasks, the pole was presented in a position that was easily reached but did require some whisking. All animals began training on dedicated training rigs. Once animals learned the task using whiskers C1-3, animals were trimmed to a single whisker (typically C2, occasionally C3). Subsequent trimming occurred every 2-3 days. To collect whisker video during behavior, mice were moved to a rig with whisker videography once they were proficient at the final version of their task. Lesions in these animals were not performed until they exhibited stable performance on the whisker videography rig, as some animals took 1-2 sessions to adjust to the new apparatus.

The behavioral task was controlled by a BPod state machine (Sanworks) and custom MATLAB software (MathWorks) running on a behavioral computer (System 76). The auditory response tone was controlled by a low-latency audio board (Bela). Lickport motion was controlled by a set of 3 motorized actuators (Zaber) and an Arduino. Licks were sensed using a custom electrical detection circuit.

### Barrel Identification

For the Ai162 X Slc17a7-Cre mice, the locations of barrels in vS1 corresponding to whiskers C1-3 was identified by measuring the GCaMP6s ΔF/F at coarse resolution (4X; 2.2 × 2.2 mm field of view) on a two-photon microscope while the whiskers were individually deflected. Mice were awake but not engaged in a task. Barrel locations in the C57/BL6J mice were identified using intrinsic signal imaging while the whiskers were individually deflected. In this case, mice previously implanted with vS1 cranial windows were anaesthetized with isoflurane. Imaging was done with a 10-bit Basler ACE camera mounted on a stereoscope and custom MATLAB software. Green illumination (Thorlabs, M530L4) was used to image the vasculature, and red illumination (Thorlabs, M625L4) was used for functional imaging. Individual whiskers were placed inside a capillary tube and stimulated with a piezo stimulator (Thorlabs, PB4NB2S). Stimulation consisted of 5 repetitions of 10 deflections, each lasting 50 ms, with a 100 ms interval in between. Deflections typically resulted in an angular displacement of 5-10° at the follicle. For each whisker, the region of high ΔR/R was mapped with respect to the vasculature, and 2-3 whiskers were mapped per mouse to confirm barrel location.

### Whisker Videography

Whisker video was acquired using custom MATLAB software from a CMOS camera (Ace-Python 500, Basler) running at 400 Hz and 640 × 352 pixels and using a telecentric lens (TitanTL, Edmund Optics). Illumination was provided by a pulsed 940 nm LED (SL162, Advanced Illumination). 7-9 s of each trial were recorded, including 1s prior to pole movement, the period when the pole was in reach, and several seconds after the pole was retracted. Data was processed on NYU’s High Performance Computing (HPC) cluster. First, candidate whiskers were detected using the Janelia Whisker Tracker(Clack et al., 2012). Next, whisker identity was refined and assessed across a single session using custom MATLAB software (Peron et al., 2015; Peron et al., 2020). Following whisker assignment, whisker curvature (κ) and angle (θ) were calculated at specific locations along the whisker’s length. As per convention, protraction yielded more positive θ values.

Change in curvature, Δκ, was calculated relative to a resting angle-dependent baseline curvature value obtained during periods when the pole was out of reach. Next, automatic touch detection was performed. Touch assignment was manually curated using a custom MATLAB user interface (Peron et al., 2015). The angle (θ) was decomposed into whisking setpoint, amplitude, and velocity via the Hilbert transform (Kleinfeld and Deschenes, 2011).

### Lesions

Cortical lesions were performed using a 1040 nm laser (Fidelity HP, Coherent) by focusing the laser at 200-300 μm depth for 10-20 s at 1-1.5 W power. Lesions were either centered on the target barrel (experimental lesions) or in visual cortex (posterior and medial relative to the target barrel; sham lesions). Lesions targeted a single site. In Ai162 X Slc17a7-Cre mice, the desired barrel was found using coarse resolution 2-photon imaging (4X), and lesion success was visually confirmed by an increase in GCaMP6s fluorescence in the target area. In C57/BL6J mice, the target barrel was found using epifluorescence imaging of the vasculature and the intrinsic imaging map as a reference. Lesion efficacy was often validated by performing an identical lesion in an Ai162 X Slc17a7 mouse immediately before lesioning of trained C57/BL6J mice. Because these mice expressed GCaMP6s, lesion extent could be calibrated based on the post-lesion fluorescence. Animals were awake and head-fixed in the behavioral apparatus during lesioning and were monitored for signs of distress or discomfort. Most animals were lesioned 5-30 minutes before the start of behavior, although a few animals were lesioned in the middle of a behavioral session.

Following behavioral training and lesioning, all animals were perfused with paraformaldehyde (4% in PBS) and post-fixed overnight. Coronal sections 70-80 μm thick were cut on a vibratome (Leica) and mounted on glass slides with VECTASHIELD antifade mounting media containing DAPI (Vector Labs). Sections were imaged using a fluorescent light microscope (VS120, Olympus).

To quantify lesion volume, we used DAPI (C57/BL6J mice) or GCAMP6s fluorescence (Ai162 X Slc17a7-Cre mice). Serial images of the lesion were collected using a fluorescent light microscope (VS120 ; Olympus). All images in which a lesion was present were registered to the Allen common coordinate framework using the SHARP-track pipeline (Shamash et al., 2018). In this way, lesions from each animal were normalized to the standard brain atlas and lesions could be compared across animals. Lesion volume was quantified by measuring the area of manually-delimited lesion borders across adjacent sections and calculating volume as: V = (A_1_ + A_2_ +. + A_n_)*t, where A_n_ is the lesion area in the final slice containing a lesion, and t is the thickness at which sections were sliced (Shih et al., 2013). For animals where some lesion slices were not recoverable, we estimated the lesion volume from the available slices. The slice with the largest lesion area was taken to be the maximum area the lesion reached across all slices. We linearly fit known volumes and maximal area across the animals for which all slices were present, and used this fit along with the maximal area to estimate lesion volume. Animals for which volume was estimated in this manner are denoted with a cross in **Table 1**.

### Immunohistochemistry

Lesions were performed as described earlier in two Ai162 X Slc17a7-Cre mice. Perfusion was performed 24 h after lesioning. 50 μm thick sections were then cut on a vibratome (Leica) and sections that included the lesion were incubated overnight with primary antibodies made in 1% bovine serum albumin and 0.05% sodium azide under continuous agitation. Alternating slices were labeled for Iba1 and glial fibrillary acidic protein (GFAP) (Podgorski and Ranganathan, 2016). Slices were washed three times and then incubated in a species appropriate secondary antibody (1:500) conjugated to Alexa Fluor 647. Slices were rinsed and mounted using antifade mounting media (Vector Laboratories). Slices were imaged using the Olympus VS120 microscope and a Leica SP5 confocal microscope.

Primary antibodies: rabbit anti-Iba1 (019-19741; Wako; 1:500 dilution), mouse monoclonal anti-GFAP (G3893; Sigma-Aldrich; 1:1,000 dilution). Secondary antibodies: goat anti-Mouse, Alexa Fluor 647 (Iba1 and HSP; Thermo Fisher A-21244), and Goat anti-Rabbit, Alexa Fluor 647 (GFAP; Thermo Fisher A-21235).

### Experimental Design and Statistical Analyses

For comparisons between distinct samples, two-tailed unpaired t-tests were used. For longitudinal comparisons within the same animals, paired t-tests were used. For correlation tests, Pearson’s correlation was used to identify a linear correlation coefficient (R) and test for significance. All statistical analysis was performed using MATLAB. Exact lists of animals used for each experiment are provided in **Table 1**.

## Results

### Mice use touch to solve a single whisker go/no-go location discrimination task

We trained water-restricted mice implanted with a cranial window over primary vibrissal somatosensory cortex (vS1) to perform a head-fixed go/no-go object location discrimination task using a single whisker (**Figure 1A**; Methods). Mice received a water reward for licking when a vertical pole was presented in a series of proximal positions (‘go’ trials). On ‘no-go’ trials, the pole appeared in a series of distal positions and licking responses resulted in a timeout. Mice became proficient at this task in 12.9 ± 2.0 days (n=10 mice; mean ± S.D.; **Figure 1B**).

**Figure 1.**
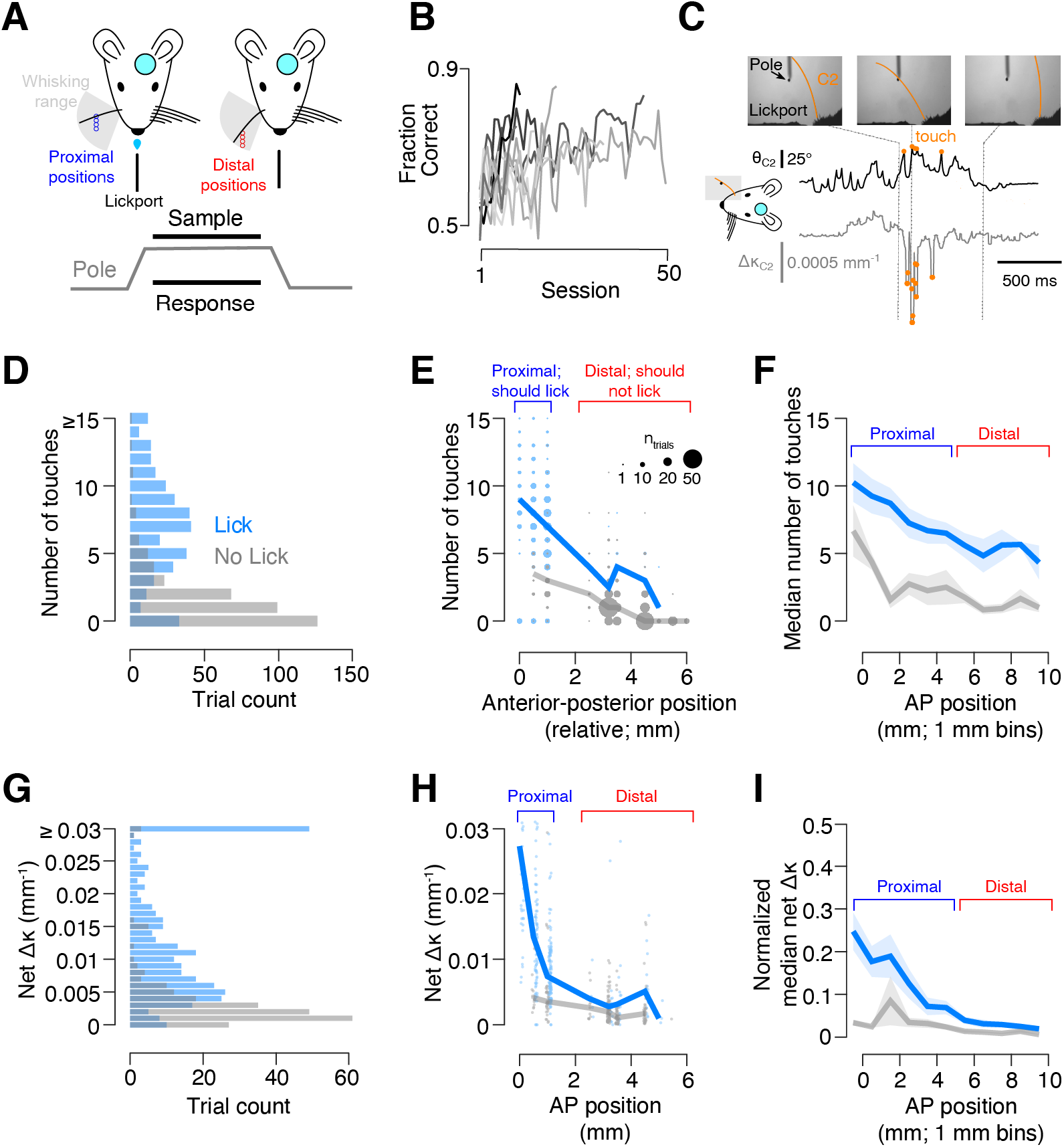
Mice use touch to solve a go/no-go object location discrimination task. **A**. Task schematic. Top, head-fixed mice with a cranial window over vS1 use a single whisker to localize a pole that appears in either posterior (blue) or anterior (red) positions. Bottom, task timing. The pole is accessible during the sample period (1-2 s) and mice must respond by licking during this period. **B**. Training progression for all go/no-go discrimination mice (n=10). **C**. Whisker videography in an example trial. Top, example frames. Bottom, whisker angle (θ, black), and change in whisker curvature (Δκ, grey, see Methods), with touches overlaid as orange circles. **D**. Number of trials with a specific touch count for the two pre-lesion behavioral sessions in an example mouse. Light blue, trials where the mouse made a lick response; grey, trials with no lick. **E**. Number of touches as a function of pole position during the 2 pre-lesion sessions for an example mouse. Thick lines, the mean for a position. Individual dots are sized to show how many trials with a given outcome, touch count, and position occurred. **F**. Number of touches across anterior-posterior pole positions for all mice averaged across the two pre-lesion sessions. Line and shaded region show mean and S.E.M. (n=9 mice). Positions across animals were aligned so that the transition between trials on which the mouse should and should not lick was at 5 mm. **G**. Number of trials with a specific touch intensity (net Δκ) across two pre-lesion behavioral sessions in an example mouse. **H**. Touch intensity (net Δκ) as a function of pole position, for an example mouse in an example session. Thick lines, the median for a position. Individual dots show individual trials and are slightly jittered along the x-axis to facilitate visibility. **I**. Touch intensity (median net Δκ, normalized to within-animal 99^th^ percentile) across anterior-posterior pole positions for all mice. Line and shaded region show mean and S.E.M. (n=9 mice). Positions across animals were aligned as in **F**.

Once mice were proficient at the task, we employed high-speed videography to assess the impact of vibrissal-object touch on behavior (Methods; **Figure 1C**). Mice were sensitive to the number of object touches, licking more often on trials with a higher number of touches (**Figure 1D**). The number of touches was position dependent, with anterior positions typically eliciting fewer touches (**Figure 1E, F**). Across all pole positions, a higher number of touches was more likely to elicit a lick response.

Vibrissal S1 touch neurons show increasing responses to higher contact forces (Peron et al., 2015). We therefore examined the impact of touch force on behavior, using the net whisker curvature change on a trial as a proxy for force acting on the whisker follicle (Birdwell et al., 2007; Pammer et al., 2013) and thus driving primary sensory afferent responses (Severson et al., 2017). Mice licked more frequently on trials with greater contact forces (**Figure 1G**). At more posterior positions, contact force was higher, and, for any given pole position, higher contact force increased the likelihood of licking (**Figure 1H, I**). Thus, both the frequency and intensity of touch impacted behavior in our task.

To ensure that mice were performing an object location discrimination task, we confirmed that mice touched reliably at both proximal and distal positions. Mice will often exhibit vibrissal foveation in object localization tasks, restricting whisker movement to the proximal positions, rather than whisking through their full whisking extent (O’Connor et al., 2010). This transforms the behavior from a discrimination to a detection task. To counteract this tendency, we used a series of proximal and distal pole positions and presented these positions at different frequencies tailored to each individual animal (**Figure 2A**). This resulted in a high number of object contacts at both proximal and distal pole positions (**Figure 2B, C**). Most mice showed performance decrements on proximal trials where no touch occurred, although a few mice showed equal performance for proximal pole positions on touch and non-touch trials (**Figure 2D**). Mice consistently performed worse on distal trials where touch occurred compared to distal trials without touch. Therefore, mice naturally tend towards a detection strategy, but our multi-position approach ensures that they continue to touch across both position ranges, necessitating that they perform a location discrimination task rather than a detection task.

**Figure 2.**
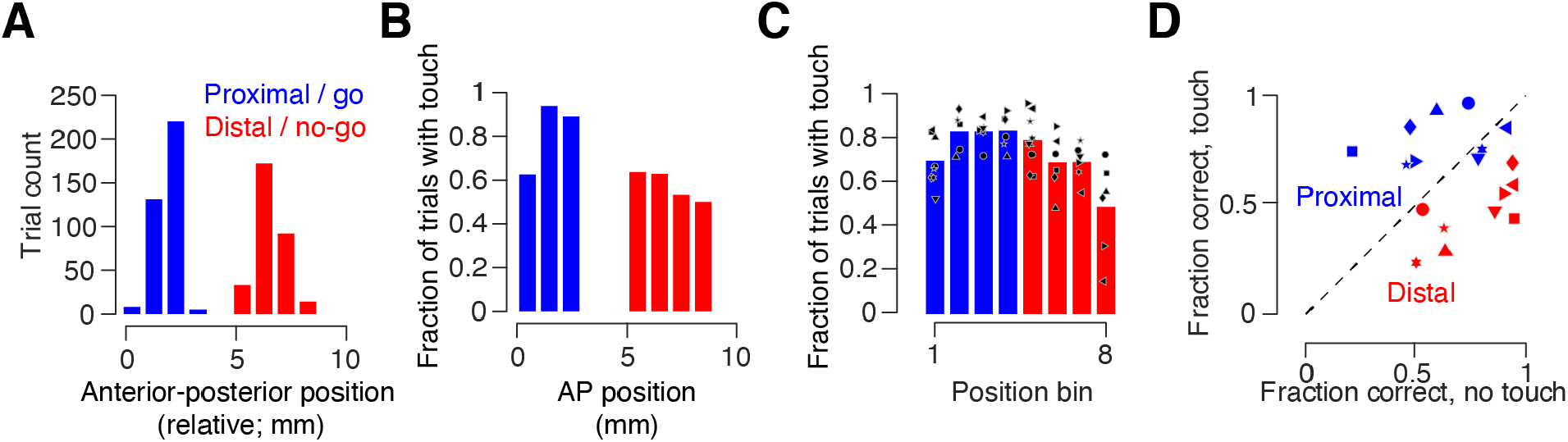
Mice touch reliably at both proximal and distal pole positions. **A**. Number of trials at each pole positions across the two pre-lesion days in an example mouse. **B**. Fraction of trials with touch on any given position for the same days as in **B. C**. Touch frequency across all pre-lesion behavioral sessions with whisker videography (Methods) across all mice (n=9). Positions were normalized across mice by binning both proximal and distal position ranges into four equal-sized bins. Symbols, individual mice. Bar, mean across mice. **D**. Modulation of performance by touch during two pre-lesion sessions. Symbols, individual mice. Blue, go/proximal trials; red, no-go/distal trials.

### Focal vS1 lesions impair performance on go/no-go location discrimination task

To determine whether individual vS1 columns are necessary for object location discrimination, we lesioned a target barrel along with portions of adjacent barrels in trained animals. We tracked performance for several days following the lesion (**Figure 3A**; Methods). Barrels were first identified using intrinsic signal imaging (**Figure 3B**) and their locations were mapped onto to the vasculature (Methods). Lesions were performed by subjecting the target barrel to prolonged femtosecond laser exposure, immediately prior to the first post-lesion behavioral session. Lesions did not require any surgery or anesthesia as they were performed on mice previously implanted with cranial windows. Lesion volumes were comparable to the volume of two barrels: 0.17 ± 0.09 mm^3^ (mean ± SEM, n = 7 mice; volume of a single barrel: 0.09 mm^3^, assuming a ∼300 μm diameter cylinder spanning 1.288 mm in cortical depth (Lefort et al., 2009); exact size: 1.9 ± 0.1 barrels, or 12,300 ± 647 neurons), and did not penetrate the white matter (**Figure 3C, D**). Lesions did not evoke distal microglial or astrocytic reactions (**Figure 3E**). Thus, our approach allowed for lesions on the scale of individual cortical columns.

**Figure 3.**
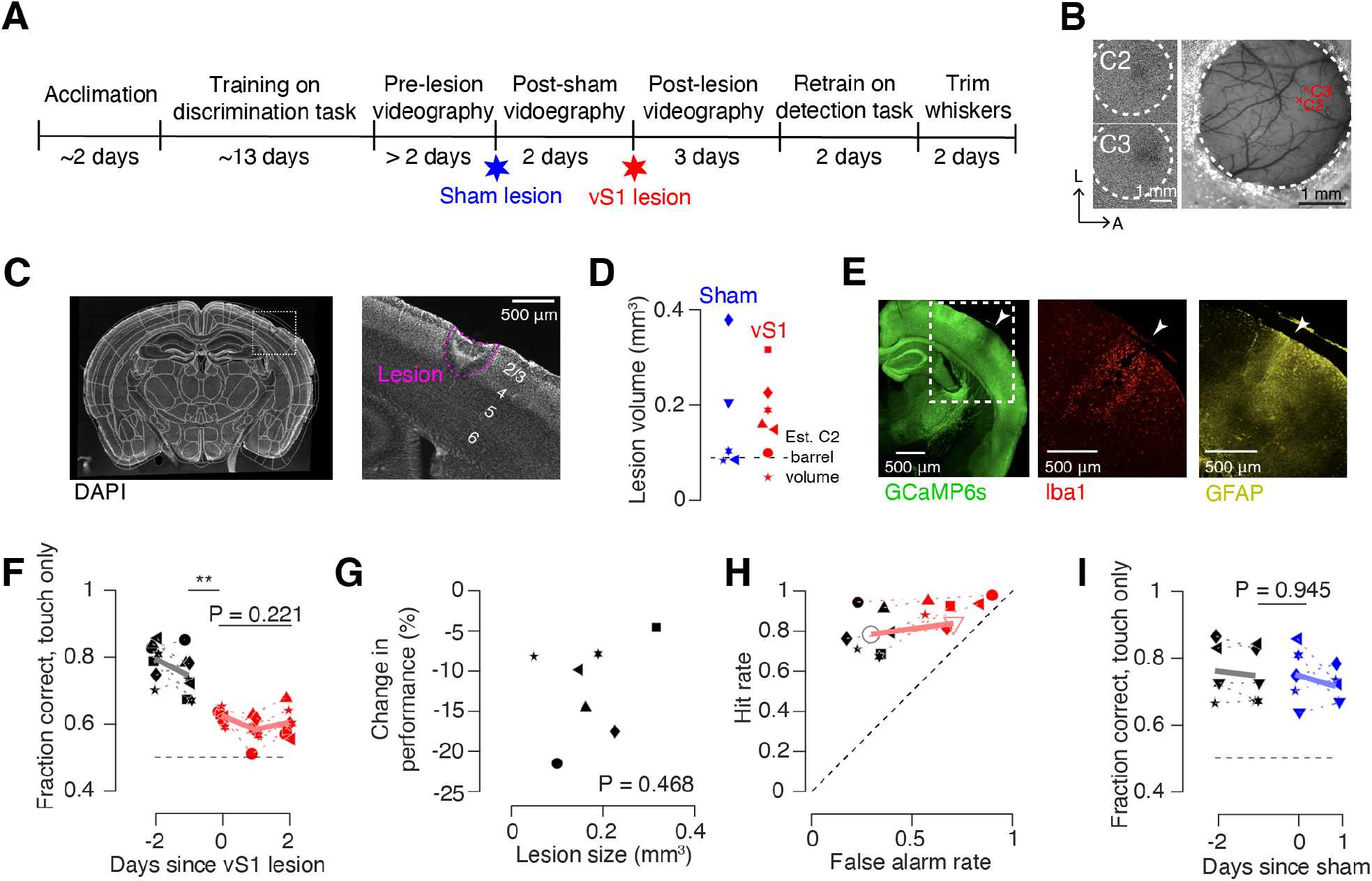
Columnar scale lesions impair performance on go/no-go discrimination task. **A**. Timeline of typical experiment. **B**. Identification of barrels. Left, intrinsic signal imaging response to individual deflection of the C2 or C3 whiskers. Dotted circle: window. Right, center of intrinsic response mapped onto the vasculature (Methods). **C**. Post-lesion histology. Example DAPI coronal section in mouse with a lesion. Left, section registered to the Allen brain atlas. Right, area around lesion, with lesion outlined in pink. **D**. Distribution of sham lesion (blue) and vS1 lesion (red) volumes. Symbols, individual mice. Dashed line, estimated C2 barrel volume. **E**. Post-lesion immunohistochemistry in an example mouse. Left, GCaMP6s. Middle, Iba1, a microglial marker. Right, sGFAP, an astrocytic marker. **F**. Performance before (black) and after (red) the lesion. Solid line, mean across mice (n=7). Symbols, individual mice. **G**. Change in performance following lesion as a function of lesion size. P-value, Pearson correlation. **H**. Change in hit and false alarm rates following lesion. Performance is for day prior to and last post lesion day. Large symbols, mean across mice. Black, pre-lesion; red, post-lesion. **I**. Performance before (black) and after (blue) sham lesion delivered to visual area. Solid line, mean across mice. **, P < 0.01.

We next examined the impact of columnar scale lesions on performance for the go/no-go location discrimination task. We recorded high-speed whisker video for at least two days prior to the lesion and three days following the lesion. To assess behavior on trials where the animal was performing object localization, we restricted our behavioral analyses to trials on which touch occurred (73.7 ± 2.7% of trials on pre-lesion sessions, 71.7 ± 4.2% of trials post-lesion, n = 7 mice). Following a lesion, performance on touch trials declined from 74.4 ± 2.5% correct to 62.4 ± 0.8% (**Figure 3F**; P = 0.002, paired t-test comparing day before to day-of lesion; n = 7 mice). We checked for recovery in the days following the lesion, but performance remained low (day 2 post-lesion: 60.3 ± 1.6%) on the third post-lesion session (P = 0.221, lesion day vs. day 2 post-lesion; n = 7). The decline in performance was not sensitive to the size of the lesion (R = 0.48, P = 0.468, Pearson correlation**; Figure 3G**). This decline was primarily due to an increase in the fraction of false alarms (from 29.2 ± 3.1 to 70.2 ± 5.6%, P < 0.001, paired t-test comparing day before to day after lesion n = 7 mice; **Figure 3H**). Sham lesions of comparable size (0.17 ± 0.13 mm^3^, n = 5 mice, P = 0.989, unpaired t-test comparing sham to vS1 lesion volumes) in visual areas posterior and medial to vS1 performed in a subset of animals did not produce behavioral effects (performance before: 74.5 ± 2.8 %, after: 74.8 ± 2.9 %; P = 0.945, n = 5; **Figure 3I**). Thus, focal vS1 lesions drive irreversible declines in performance on object location discrimination tasks.

The increase in false alarm rate on no-go (distal position) trials suggested that post-lesion, mice adopted a detection strategy, licking upon object contact regardless of object position and withholding licking when touch did not occur. To test this hypothesis, following the first three post-lesion days, mice were transitioned to a detection task in which the distal pole positions were completely out of reach (**Figure 4A**). Upon moving lesioned mice to this task, their performance improved from 58.4 ± 1.8% to 88.9± 2.5% (**Figure 4B**; P < 0.001, paired t-test comparing last post-lesion discrimination day vs. first detection day; n=7 mice). This improvement was mostly due to a decline in false alarm rate (from 71.6 ± 5.1% to 10.0 ± 2.7%, P < 0.001, n = 7 mice; **Figure 4C**), implying that lesions had not impaired the capacity of mice to perform the go/no-go task at high levels but instead had interfered with sensory input. In all mice, performance on this detection task exceeded the performance on the final pre-lesion discrimination task day (P < 0.001, n = 7) implying that mice performing the discrimination task are fundamentally perceptually limited relative to mice performing the detection task.

**Figure 4.**
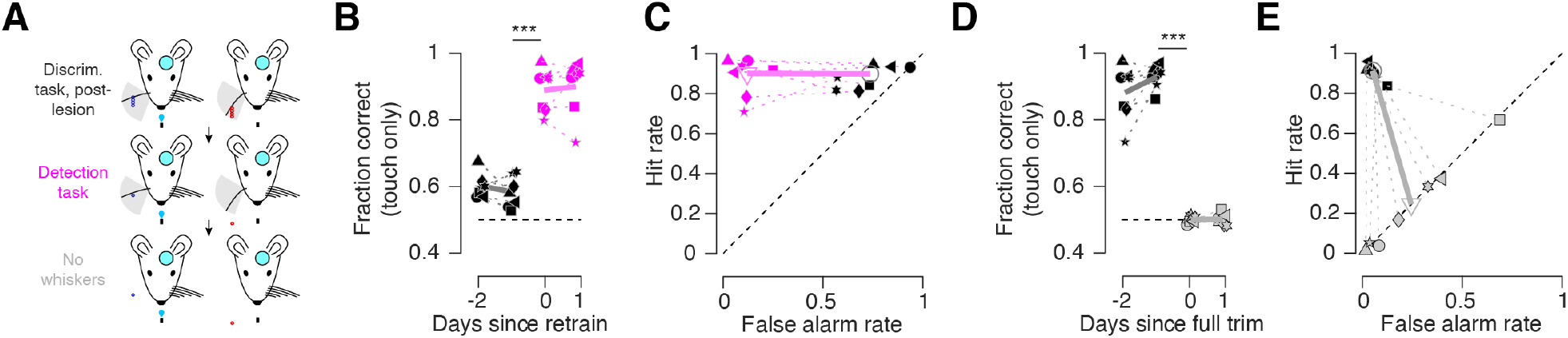
Lesioned animal detection task and post-trim performance. **A**. Transition from post-lesion discrimination to detection, and then to full whisker trimming. **B**. Fraction correct before (black) and after (magenta) transition to detection task. Symbols, individual mice. Thick line, average across mice. **C**. False alarm and hit rate before (black) and after (magenta) transition to detection task. Open symbols, average across mice. **D**. Fraction correct before (black) and after (grey) removal of whisker. **E**. False alarm and hit rate before (black) and after (grey) whisker removal. ***, P < 0.001.

If mice had indeed adopted a lick-upon-touch strategy following the lesion, mice experiencing no touch should withhold licking. To evaluate this possibility, we cut the remaining whisker after two days of the detection task and examined the impact on behavior. Performance dropped to chance (performance before trimming: 86.5 ± 4.6%, after: 50.0 ± 0.5%; P < 0.001, n = 7; **Figure 4D**). All mice adopted a constant lick rate, as evidenced by a matched hit and false alarm rate (**Figure 4E**), consistent with a guessing strategy. Most mice licked on less than 25% of trials, substantially less than before trimming, despite their being highly motivated to do so. This is consistent with a lick-upon-touch strategy.

Thus, columnar scale vS1 lesions disrupted performance on an object location discrimination task. Mice adopted a lick-upon-touch strategy following the lesion, as demonstrated by their ability to learn a detection task and near-cessation of licking following whisker trimming.

### Focal vS1 lesions do not impact vibrissal kinematics

Area scale lesions and optogenetic silencing of vS1 alter vibrissal kinematics (Hong et al., 2018), with mice exhibiting reduced whisking vigor and lower touch intensities. Do columnar scale lesions drive shifts in vibrissal kinematics? To address this question, we compared kinematics before and after lesions (**Figure 5A**). In mice performing the go/no-go location discrimination task, columnar scale lesions did not show any effect on the kinematic parameters we measured. Mice made 3.6 ± 1.4 touches per trial before the lesion, and 3.9 ± 2.4 touches per trial after (**Figure 5B**; P = 0.872, paired t-test, n = 7 mice). The intensity of the touches also did not change, quantified using the net curvature change per trial (net Δκ, **Figure 5C**; before lesion: 0.014 ± 0.006 mm^-^1^; after lesion: 0.023 ± 0.013 mm^-^1^, P = 0.079; Methods). Peak curvature change was not impacted by lesions (**Figure 5D**; before lesion: 0.0011 ± 0.0006 mm^-^1^; after lesion: 0.0013 ± 0.0005 mm^-^1^, P = 0.395). Whisking intensity also remained constant (**Figure 5E-G**; Methods), quantified using peak whisking amplitude (before lesion: 3.5 ± 2.1°, after lesion: 3.4 ± 1.5°, P = 0.780), peak setpoint (before lesion: 4.1 ± 6.4° after lesion: 6.7 ± 5.9°, P = 0.325), and peak velocity (before lesion: 332.6 ± 177.8°/s after lesion: 291.54 ± 74.3°/s, P = 0.579). Thus, whisking kinematics were not altered by columnar scale lesions, implying that observed behavioral changes were not due to changes in afferent input.

**Figure 5.**
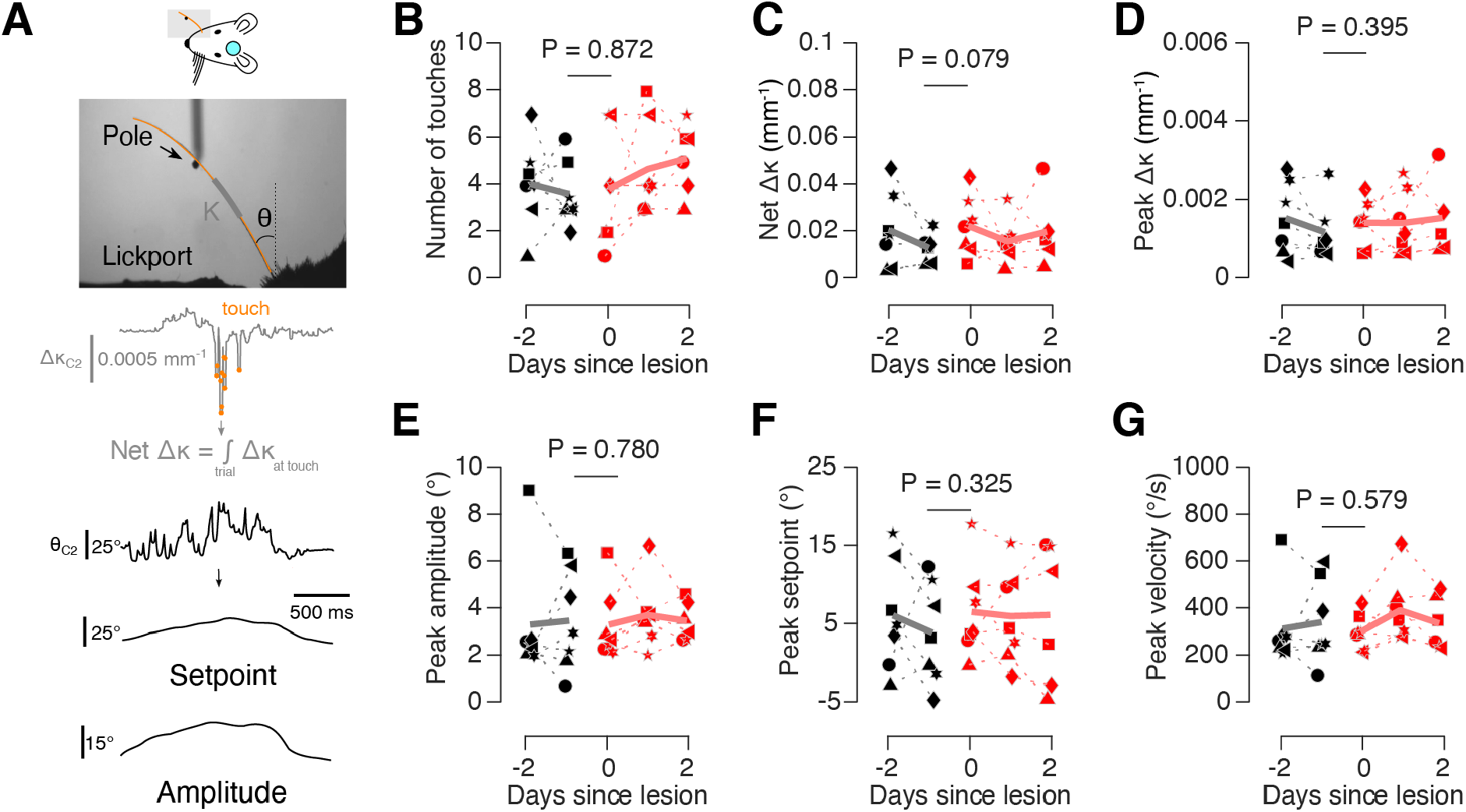
Lesions do not impact vibrissal kinematics. **A**. Kinematic variables measured. Top, example whisker video frame, showing curvature (Δκ, grey) and angle (θ, black) measurement. Middle, change in whisker curvature (Δκ, grey), with touches overlaid as red circles, and calculation of net Δκ. Bottom, whisker angle over time (θ, black), with whisking setpoint and amplitude obtained from Hilbert transform (Methods).**B**. Number of touches per trial prior to lick response before (black) and after (red) lesioning in mice performing the go/no-go discrimination task. Thick line, cross-animal mean. P-value provided for paired t-test comparing pre-lesion day to day of lesion, n=7 mice. **C**. Mean net curvature change (net Δκ), a measure of total touch force over a trial, up to moment of first lick before (black) and after (red) lesion. **D**. Peak curvature change (Δκ), a measure for maximum force exerted during touch, before (black) and after (red) lesion. The ‘peak’ was the 99^th^ percentile of values over epochs during which the mouse was touching the pole. **E**. Peak whisking amplitude (Methods) before (black) and after (red) lesion. The peak was the 99^th^ percentile of values over epochs during which the pole was in reach but prior to the first touch. **F**. Peak whisking setpoint (Methods) before (black) and after (red) lesion. The peak was the 99^th^ percentile of values over epochs during which the pole was in reach but prior to the first touch. **G**. Peak whisker velocity before (black) and after (red) lesion. The peak was the 99^th^ percentile of values over epochs during which the pole was in reach but prior to the first touch.

### Focal lesions do not interfere with simple or complex vibrissal object detection tasks

Mice with vS1 lesions show degraded performance on an object location discrimination task but attain high performance on an object detection task (**Figure 4B**). We therefore asked whether columnar scale lesions to vS1 would perturb performance in mice trained only on a detection task. Following area scale vS1 lesions, mice exhibit temporary declines in performance on a go/no-go detection task (Hong et al., 2018). We therefore trained a separate cohort of mice on a go/no-go vibrissal object detection task (**Figure 6A**). Mice learned this task more readily than the discrimination task, reaching above-threshold performance in 5.0 ± 0.5 days (**Figure 6B**; n = 6 mice; P < 0.001, t-test comparing detection and discrimination cohorts). We observed no effect on performance following the lesion, despite lesions taking place within an hour of the start of the post-lesion behavioral session (performance before lesion: 80.9 ± 1.7%, after: 75.7 ± 2.2%; P = 0.054, n = 6 mice; **Figure 6C**). Lesion size (0.52 ± 0.29 mm^3^, n = 6 mice) did not correlate with the effect on performance (**Figure 6D**; P = 0.308, Pearson correlation). We confirmed that this was a whisker-dependent behavior by trimming the whisker after 3 post-lesion days. Following whisker trimming, mice performed at chance (performance before trimming: 83.9 ± 2.1%, after: 50.3 ± 1.4%; P < 0.001, n = 5; **Figure 6E**). Thus, simple touch detection behaviors are not vS1 dependent, and previously observed declines in performance in such tasks are likely due to the extensive size of those lesions.

**Figure 6.**
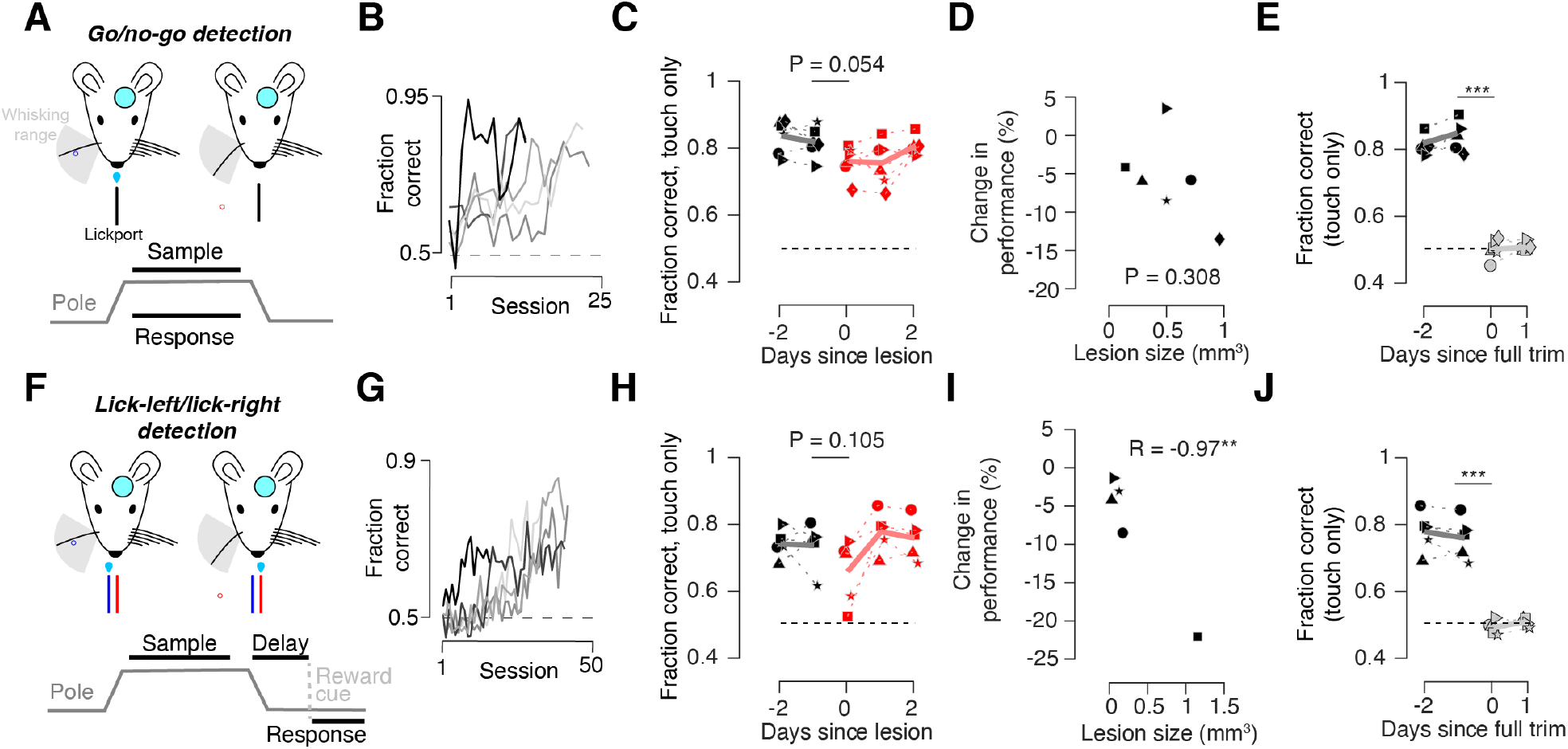
Lesions do not impair performance on detection tasks. **A**. Task schematic for go/no-go detection task. Top, mice with a cranial window use a single whisker to localize a pole that appears in either a posterior (blue) or out of reach, anterior (red) position. Bottom, task timing. **B**. Training progression for all go/no-go detection mice (n=6). **C**. Performance before (black) and after (red) the lesion. Solid line, mean across mice (n=6). P-value provided for paired t-test comparing pre-lesion day to day of lesion. **D**. Change in performance as a function of lesion size. P-value indicated for Pearson correlation. **E**. Fraction correct before (black) and after (grey) removal of last remaining whisker. **F**. Task schematic for lick-left/lick-right (2 lickport) detection task. Top, mice with a cranial window use a single whisker to localize a pole that appears in either a posterior (blue) or out of reach, anterior (red) position, and lick one lickport to report touch, and the other to report no touch. Bottom, task timing. Animals respond after a short delay following the sample period upon reward cue. **G**. Training progression for all lick-left/lick-right detection mice (n=5). **H**. Performance before (black) and after (red) the lesion. Solid line, mean across mice (n=5). **I**. Change in performance as a function of lesion size. P-value indicated for Pearson correlation. **J**. Fraction correct before (black) and after (grey) removal of last remaining whisker. **, P < 0.01; ***, P < 0.001.

Finally, because tasks with higher cognitive load are believed to be more dependent on cortical activity than simpler tasks (Stuttgen and Schwarz, 2018), we examined the impact of columnar scale lesions on a more complex detection behavior. We trained mice to perform a lick-left/lick-right touch detection behavior that required mice to report touch by licking one of two lickports and the lack of touch by licking the other lickport (**Figure 6F**). In contrast to the simpler go/no-go behaviors, this task required mice to withhold licking until 0.5 s after the pole became inaccessible, at which point an auditory cue indicated to the animal that it was time to respond. Thus, this task had both two response contingencies and a working memory component. Mice learned this task more slowly than the go/no-go detection task, reaching above-threshold performance in 23.4 ± 3.9 days (**Figure 6G**; n = 5 mice; P < 0.001, t-test comparing go/no-go and lick-left/lick-right detection cohorts). Despite the greater difficulty in learning this task, mice again showed no performance decline following a columnar scale lesion in vS1 (performance before lesion: 73.2 ± 3.2%, after: 65.4 ± 4.4%; P = 0.105, n = 5; **Figure 6H**). Lesion size (0.31 ± 0.47 mm^3^, n = 5 mice) predicted the effect on performance (**Figure 6I**; R = -0.97, P = 0.007, Pearson correlation). Performance fell to chance following whisker trimming (performance before trimming: 75.5 ± 2.8%, after: 48.8 ± 0.9%; p < 0.001, n = 5; **Figure 6J**). Thus, even relatively complex vibrissal touch detection behaviors are not vS1-dependent.

## Discussion

We tested the behavioral role of vS1 columns in a range of vibrissa-dependent tasks and found that lesioning a barrel in vS1 impacts performance when the mouse must report whether it touched a proximal or distal object. Lesions had no impact when the mouse had to report whether it touched an object or not. The deficit in discrimination task performance did not recover after several days following the lesion. In contrast, detection performance was not impacted by columnar scale lesions even when the task was more complex. Thus, neurons within the principal whisker column in vS1 are necessary for vibrissal object location discrimination.

Vibrissal S1 has been implicated in many whisker behaviors using a range of inactivation approaches. These approaches, both transient and permanent, inactivate nearly all of vS1 (but see (Shih et al., 2013)). Spatially extensive inactivation can impact adjacent structures, which can lead to misattribution of behavioral role. For example, lesions of vS1 that extend to striatum can produce permanent performance degradation in touch detection tasks where lesions confined to vS1 produce behavioral recovery (Hong et al., 2018). Area-scale lesions of vS1 have led to degraded performance in tasks requiring discrimination of aperture size (Krupa et al., 2001), object location (O’Connor et al., 2010), distance (Hutson and Masterton, 1986), and texture (Guic-Robles et al., 1992). In contrast, object touch detection tasks show either no sensitivity to lesions (Hutson and Masterton, 1986) or transient sensitivity (Hong et al., 2018), with rapid recovery back to pre-lesion performance. Local pharmacological inactivation degrades performance in a single-vibrissa object location discrimination task (O’Connor et al., 2010) as well as tasks that require comparison of stimuli across two sides of the head (Shuler et al., 2002; Miyashita and Feldman, 2013). Optogenetic silencing of vS1 interferes with object location discrimination (Guo et al., 2014b) and object detection (Hong et al., 2018) tasks. Spatial dissociation of function in auditory cortex demonstrates that specific regions subserve specific functions (Lomber and Malhotra, 2008). Consequently, whether these tasks require large portions of vS1, surrounding areas that are likely impacted by past inactivation approaches, or just the relevant whisker barrel(s) remains unclear. Our work suggests relatively small populations of neurons likely underpin many behavioral functions.

Gap crossing with a single whisker, which is impeded following large scale vS1 lesions (Hutson and Masterton, 1986), is susceptible to microstrokes that produce lesions on the scale of a single barrel (Shih et al., 2013). These strokes are generated by occluding blood vessels either via laser illumination of a sensitizer or via amplified laser pulses. Columnar scale lesions are comparable in size to microstrokes, but are produced without the need for targeting blood vessels or the use of special reagents. Both approaches demonstrate that relatively small (10,000-20,000) populations of neurons can contribute substantially to cortically dependent behaviors, raising the intriguing possibility that an even smaller subset of these neurons may act as a bottleneck in the generation of perception.

What is the minimal set of neurons needed for the discrimination of object location in vS1? In L2/3, approximately 10% of neurons show robust responses to touch (Crochet et al., 2011; Peron et al., 2015). L4 excitatory neurons produce fewer spikes, on average, than L2/3 neurons and L5 neurons produce comparable spike counts (Yu et al., 2019). This suggests that a sparse minority of the ∼6,000 excitatory neurons in a barrel (Lefort et al., 2009) contribute to the touch response. It is thus possible that several hundred neurons act as a perceptual bottleneck in vS1 for single whisker object location discrimination. In contrast to columnar scale lesions, however, cellular resolution lesions in vS1 targeting tens of neurons have not produced behavioral effects (Peron et al., 2020). This result is unsurprising given that those animals were performing a detection task, which we have shown to not be dependent on activity from vS1. Cellular resolution lesion experiments in mice performing discrimination tasks will thus be crucial in determining whether small populations of neurons constitute a perceptual bottleneck in vS1.

Artificially adding activity to small numbers of cortical neurons can produce behavioral effects, though this does not imply that naturalistic behaviors depend on such small numbers of neurons. In vS1, mice can readily detect the activation of arbitrary groups of tens of neurons (Dalgleish et al., 2020). Some neurons can drive behavioral report individually (Houweling and Brecht, 2008), with touch-sensitive excitatory neurons showing the most consistent ability to drive single neuron perceptual report (Tanke et al., 2018). In mouse primary visual cortex (Carrillo-Reid et al., 2019; Marshel et al., 2019), activating a handful of neurons can perturb discrimination between vertical and horizontal gratings. These experiments do not, however, demonstrate that naturalistic percepts only require tens of neurons. Strong cortical recurrence will amplify artificially introduced spikes (London et al., 2010), leaving open the possibility that under naturalistic input, many more than the neurons perturbed in these experiments are needed for behavior. For example, in tongue premotor cortex, activating ∼10 neurons can perturb the chosen licking direction (Daie et al., 2019), but silencing does not exert an effect unless hundreds of thousands of neurons are inactivated (Guo et al., 2014a). Moreover, adding activity to hundreds of L4 vS1 neurons only partially increases the report of touch (O’Connor et al., 2013). Thus, while gain-of-function experiments are crucial to our understanding of the neural basis of perception, loss-of-function experiments are needed to establish that a given population plays a role in a naturalistic behavior. Our experiments establish a ceiling of ∼10,000 excitatory neurons as a perceptual bottleneck for discrimination of naturalistic touch.

Transient perturbations of a target structure can produce behavioral effects due to temporary disruptions in downstream activity. In some cases, transient inactivation of a structure can thus lead to a behavioral deficit while removal of that structure does not influence behavior or leads to rapid recovery (Otchy et al., 2015; Hong et al., 2018). Identifying neurons that underpin perception ultimately requires identifying structures whose removal permanently perturbs behavior. Finding these structures requires permanent inactivation followed by longitudinal observation. Lesions are well suited to this task. Our work suggests vS1 barrels are chronically necessary in object location discrimination, as the behavioral impact of focal lesions persisted for several days (Wolff and Olveczky, 2018).

Primary sensory cortices are thought to be more important for behaviors involving more complex stimuli. Object location discrimination requires cortex, whereas detection does not. In the auditory cortex, many discrimination behaviors require cortex (Lomber and Malhotra, 2008; Slonina et al., 2022). At the same time, simple auditory features such as frequency can be discriminated even without cortex (Ohl et al., 1999). Lesions to V1 in humans result in ‘blindsight’ – the ability to perform certain vision-dependent tasks despite the lack of conscious visual awareness (Leopold, 2012). Complex visual tasks, however, can no longer be performed. In vS1, performance on object detection tasks recovers or does not degrade at all after inactivation (Hutson and Masterton, 1986; Hong et al., 2018), while object location discrimination (O’Connor et al., 2010) or gap crossing (Hutson and Masterton, 1986) task performance is affected. Thus, tasks involving more complex stimuli are often cortically dependent. Our work suggests these complex stimuli can be represented by relatively small neural populations.

In sum, we find that barrel columns contribute to single whisker vibrissal object localization. At the same time, vibrissal object detection is not vS1 column-dependent. Thus, small (∼10,000) populations of primary sensory cortical neurons can contribute to specific behaviors.

## Acknowledgments

We thank Dan Sanes for comments on the manuscript, and Dora Angelaki, Rob Froemke, and Bettina Voelcker for discussion. This work was supported by the Whitehall Foundation and the National Institutes of Health (R01NS117536). The authors declare no competing financial interests.

